# Full-length optic nerve regeneration in the absence of genetic manipulations

**DOI:** 10.1101/2022.08.01.502242

**Authors:** Qian Feng, Kimberly Wong, Larry I. Benowitz

## Abstract

The inability of mature retinal ganglion cells (RGCs) to regenerate axons after optic nerve injury can be partially reversed by manipulating cell-autonomous and/or -non-autonomous factors, among which are neuroimmune interactions. We report here that preconditioning resulting from a mild lens injury (conditioning LI, cLI) prior to optic nerve damage induces far greater axon regeneration than LI or the pro-inflammatory agent zymosan after nerve injury or preconditioning with Zymosan. Unlike other instances of immune-supported regeneration, cLI is unaltered by depleting mature neutrophils, T cells or blocking receptors for identified inflammation-associated growth factors (Oncomodulin, SDF1, CCL5), and is only partially diminished by suppressing peripheral monocyte recruitment. Repeated LI leads to full-length optic nerve regeneration, and pharmacological removal of local resident macrophages with the colony stimulating factor 1 receptor (CSF-1R) inhibitor PLX5622 enables some axons to re-innervate the brain in just 6 weeks. Thus, cell non-autonomous interventions not involving genetic manipulations can induce high levels of optic nerve regeneration, paving the way to uncover potent, translatable therapeutic targets for CNS repair.

## Introduction

Central nervous system (CNS) damage in adult mammals can lead to irreversible and untreatable disabilities due to neuronal death and the inability of most CNS neurons to regenerate lengthy axons. Following optic nerve damage, retinal ganglion cells (RGCs), the projection neurons of the eye, cannot extend axons beyond the injury site and soon begin to die. Intraocular inflammation induced by zymosan, lens injury (LI), or certain other manipulations partially protect RGC from dying and stimulate axon regeneration through the action of oncomodulin (Ocm), SDF-1, and other factors expressed by infiltrative myeloid cells in the mouse optic nerve crush (ONC) model^1–9^.

Conditioning refers to a phenomenon in which the application of stress before injury leads to subsequent tissue resilience or tolerance. Conditioning-induced neuroprotection in the CNS can be achieved by ischemia/hypoxia and inflammatory mediators, and conditioning-induced enhancement of axon regeneration in mammals can occur in the peripheral nervous system (PNS)^10–13^. Conditioning peripheral nerve (PN) injury potentiates the ability of dorsal root ganglion (DRG) neurons to regenerate their peripheral axon branches after a second injury and their central axon branches after spinal cord injury^14,15^. This phenomenon is primarily driven by infiltrative myeloid cells^16–19^ and represents one of the strongest known instances of spinal cord regeneration^20^. Here, we investigated whether inflammatory preconditioning would enable robust axon regeneration in the optic nerve.

## Results

### Pre-conditioning by lens injury induces robust optic nerve regeneration

Based on the role of inflammation in the PN preconditioning effect and our incidental observation that intraocular injections prior to ONC that inadvertently injure the lens result in stronger regeneration than expected, we evaluated the pro-regenerative effects of LI and intraocular zymosan before vs. after ONC in 129S1 mice (Fig. 1A). The controlled, mild LI used here disrupted the lens capsule and cortex locally without causing global cataract formation (Fig. S1A). As expected, LI or Zymosan administered immediately after ONC (LI post-ONC, zymo post-ONC, respectively) induced moderate levels of optic nerve regeneration (Fig. 1B, C), and Zymosan injected 2 weeks before ONC, like PBS controls, had almost no effect. In contrast, LI two weeks prior to ONC (cLI) nearly tripled the level of optic nerve regeneration compared to LI- or zymo-post-ONC and increased RGC survival (Fig. 1B, D). LI one week prior to ONC was far less effective than at two weeks beforehand (Fig. S1B), but cLI at 14 days and again at 3 days before ONC (1^st^ and 2^nd^ LI) greatly increased axon numbers compared to a single cLI (Fig. 1E, S1C). Two additional LI events 14 and 28 days post-ONC enabled many axons to regenerate the full-length of the optic nerve and across the chiasm 6 weeks after ONC (Fig. 1F, G) though without reaching central target areas.

**Figure 1.**
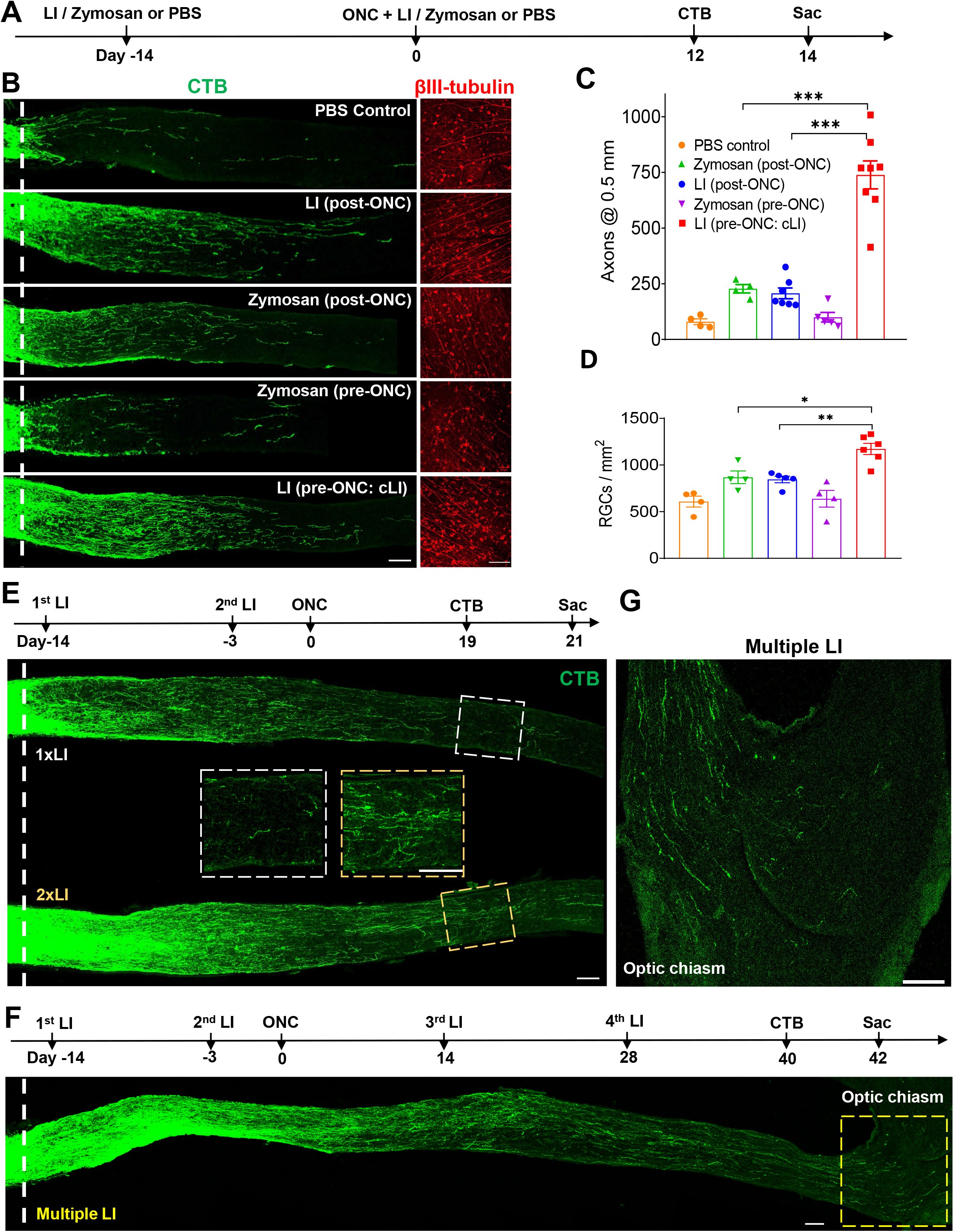
Robust optic nerve regeneration by conditioning LI (cLI). A. Experimental timeline. Lens injury, zymosan or PBS was introduced 14 days before or immediately after optic nerve crush (ONC). Mice were euthanized 14 days later. B. Left panel: representative longitudinal sections through the optic nerve showing CTB-labeled regenerating axons 14 days post-ONC. Pre-conditioning by LI before crush (LI pre-ONC: cLI) resulted in many more regenerating axons than LI post-ONC, zymosan pre-ONC, or zymosan post-ONC. *White line* indicates crush site. Right panel: representative wholemounted retinas showing βIII-tubulin^+^ RGCs. C. Quantitation of regenerating axons 0.5 mm from crush site 2 weeks after ONC (one-way ANOVA followed by Tukey’s multiple-comparisons test; p < 0.0001, n = 4 or 5 mice in each group. D. Quantitation of βIII-tubulin^+^ cells (surviving RGCs) in (B) (one-way ANOVA followed by Tukey’s multiple comparisons test; LI (pre-ONC: cLI) vs. Zymosan (post-ONC) p= 0.0190; vs. LI (post-ONC) p = 0.0064; n = 4-5 mice in each group; 7–8 fields were analyzed for each retina). E. *Top*: experimental timeline. 1^st^ LI was introduced 14 days before and 2^nd^ LI 3 days before ONC. Mice were euthanized 3 weeks later. *Bottom:* representative longitudinal sections through the optic nerve showing axon regeneration 3 weeks after ONC in mice treated with 1XLI (1^st^) vs. 2XLI (1^st^ and 2^nd^). The boxes in the center show magnified images of axons 2 mm distal to the crush site. *Dashed white line* shows crush site. Scale bar, 100 μm. Quantitation of results is shown in Fig. S1C. F. *Top*: experimental timeline. 1^st^ LI was introduced 14 days before ONC, 2^nd^ LI 3 days before, and 3^rd^ and 4^th^ LI 14 and 28 days post-ONC. Mice were euthanized 6 weeks after ONC. *Bottom*: representative longitudinal sections through the optic nerve showing fulllength optic nerve regeneration. *White line* indicates crush site. *Yellow dashed box* indicates optic chiasm. G. Enlarged image showing many regenerating axons entering optic chiasm in the yellow dashed box in (F).

### Role of major immune cell populations

Previous studies have shown that microglia are irrelevant for optic nerve regeneration induced by LI post-ONC^21^ and that zymosan-induced regeneration is mediated primarily by infiltrative neutrophils and macrophages^3–5^. To deplete resident macrophages, including microglia, perivascular macrophages and border-associated macrophages, mice were fed chow containing the Csf1r inhibitor PLX5622 starting four weeks prior to ONC (*i.e*., 2 weeks before cLI or sham surgery) whereas control mice were fed chow without drug (Fig. 2A). In mice fed standard chow, cLI doubled the number of retinal Iba1^+^ cells compared to sham-surgery controls. PLX5622 almost eliminated these cells throughout the retina after ONC and sham surgery, although after cLI, a small population of Iba1+ cells remained (Fig. S2A, C). On the other hand, in the optic nerve, a population of CD68^+^ macrophages persisted with PLX5622 treatment after either sham surgery or cLI, particularly around the crush site (Fig. S2D). Although PLX treatment alone did not promote regeneration, PLX combined with cLI nearly doubled the number of axons extending the full length of the optic nerve (≥ 4 mm beyond the injury site: Fig. 2B, C) compared to cLI alone at 4 weeks after ONC without affecting RGC survival (Fig. S2A, B).

**Figure 2.**
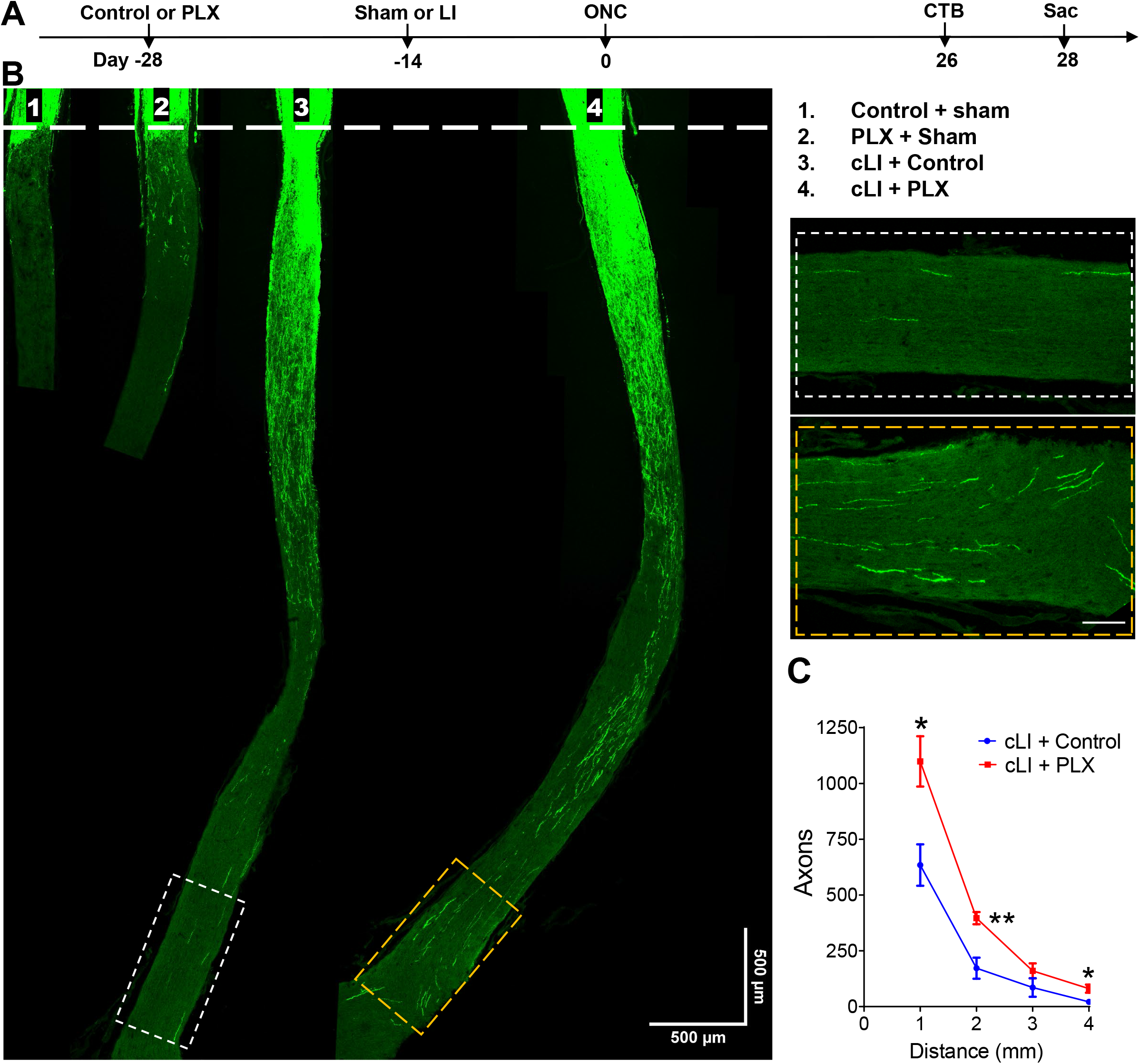
Removing naïve resident macrophages further enhances cLI-induced regeneration. A. Experimental timeline. Mice were fed with PLX or control chow from 14 days prior to LI or sham surgery until time of euthanasia. LI or sham surgery was performed 14 days before ONC, and mice were euthanized 4 weeks later. B. Representative longitudinal sections through the optic nerve showing CTB-labeled regenerating axons day 28 post-ONC. Note increase in regeneration with addition of PLX treatment. Images on right show regenerating axons near optic chiasm in the *white* and *yellow dashed boxes. White line* indicates the crush sites. Scale bar, 500 μm. C. Quantitation of regenerating axons at multiple distances from crush sites in (B) (unpaired t-test, p = 0.0131 at 1 mm; 0.0035 at 2 mm, 0.200 at 3 mm and 0.018 at 4 mm. n = 5 mice per group).

Despite being a highly selective brain-penetrant CSF1R inhibitor, PLX5622 is reported to also suppress peripheral myeloid cell populations in C57/BL6 mice^22^. However, we found that PLX treatment for 4 weeks in 129S1 mice increased the percentage of circulating monocytes (CD11b^+^Ly6C^+^Ly6G^low^ cells) in the blood without affecting the overall myeloid cell population (CD11b^+^ cells: Fig. S2E, F).

To investigate the role of peripheral monocytes, we suppressed the infiltration of these cells using mice genetically deficient in CCR2, a C-C chemokine receptor that is important for leukocyte egress from bone marrow and that is highly expressed in blood monocytes but absent in resident CNS macrophages^23^. CCR2^RFP/RFP^ mice were used as CCR2-deficient reporter mice, which had a 84% reduction in circulating CCR2^+^ monocytes (RFP^+^ cells in blood)^23^ and a 80% reduction in vitreous infiltrating CCR2^+^ monocytes (RFP^+^ cells in the vitreous chamber) compared to control heterozygous CCR2^RFP/+^ control mice following cLI and ONC (Fig. 3A, S3A, B). CCR2^RFP/RFP^ mice showed a 38% decline in cLI-induced regenerating axons 2 weeks after ONC (Fig. 3B, C) compared to heterozygous CCR2^RFP/+^ controls without diminishing RGC survival (Fig. S3C, D). Taken together, these data demonstrate that removing naïve resident macrophages with PLX5622 further enhances cLI-induced optic nerve regeneration, whereas blocking CCR2^+^ monocytes through CCR2 deficiency partially decreases regeneration. Neither of these treatments affects RGC survival.

**Figure 3.**
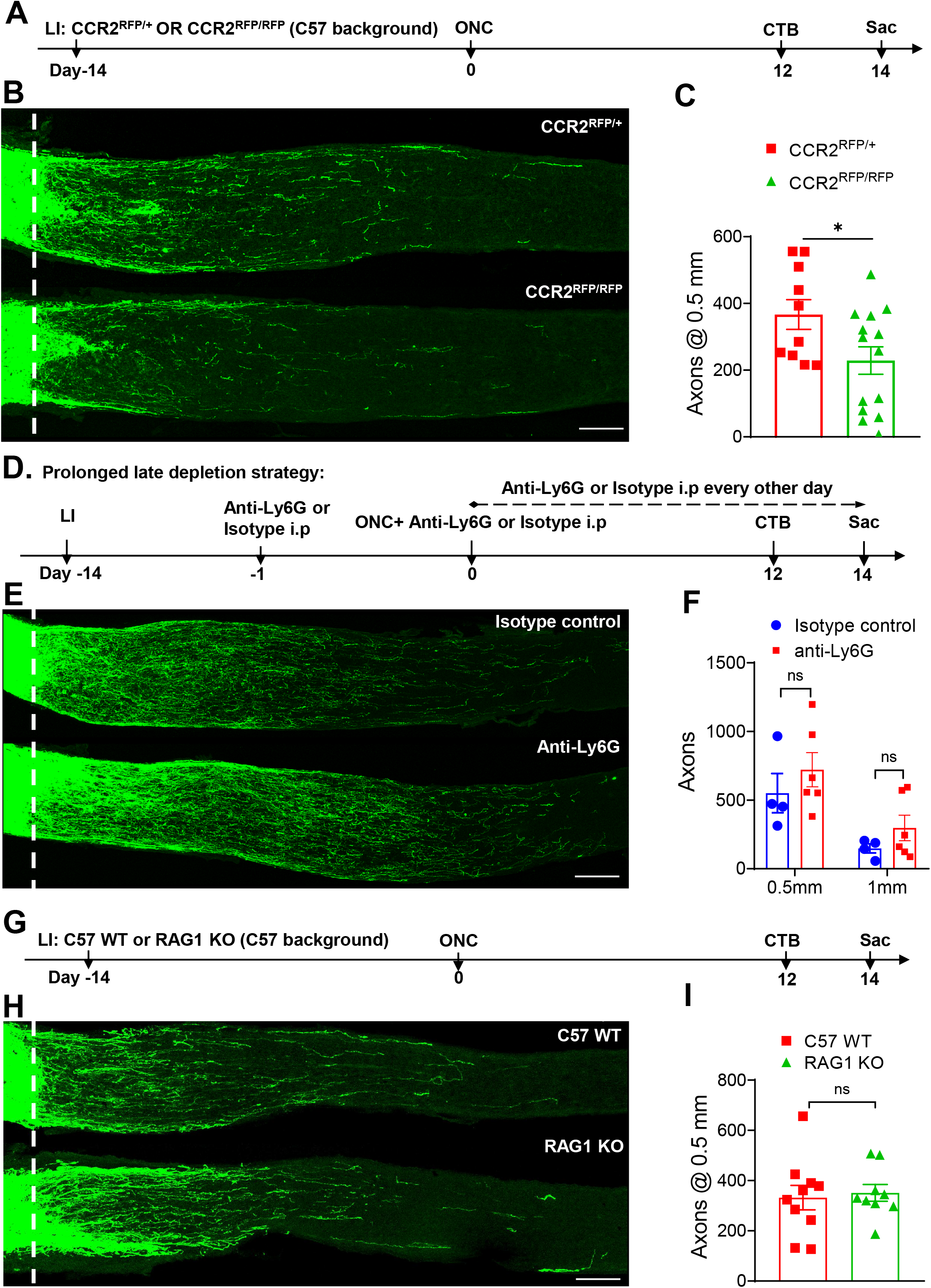
Blocking CCR2^+^ monocytes partially suppresses cLI-induced regeneration, whereas mature neutrophils and T cells are dispensable. A. Experimental timeline. LI was introduced to CCR2^RFP/+^ or CCR2^RFP/RFP^ mice 14 days before ONC. Mice were euthanized 14 days later. B. Representative longitudinal sections through the optic nerve showing decreased regeneration in CCR2^RFP/RFP^ *vs*. CCR2^RFP/+^mice. *White line:* crush site. Scale bar, 100 μm. C. Quantitation of regenerating axons (unpaired t-test, p = 0.0357; n = 6 or 7 mice per group). D. Experimental timeline for prolonged late depletion strategy. E. Representative longitudinal sections through the optic nerve show similar levels of regenerating axons in isotype control and neutrophil-depleted mice. *White line:* crush site. Scale bar, 100 μm. F. Quantitation of regenerating axons 0.5 mm and 1 mm from crush sites in (F) (unpaired t-test, p = 0.402 at 0.5 mm, p = 0.246 at 1 mm, n = 4 or 5 mice per group). G. Experimental timeline. cLI was introduced to C57/BL6J WT mice or RAG1 KO mice 14 days before ONC and mice were euthanized 14 days later. H. Longitudinal sections through the optic nerve showing similar levels of regenerating axons in C57 WTand RAG1 KO mice. *White line*: crush site. Scale bar, 100 μm. I. Quantitation of regenerating axons 0.5 mm from crush site in *H* (unpaired t-test, p = 0.758, n = 5 or 6 mice per group).

We also tested multiple strategies to immune-deplete CD11b^+^ Ly6G^+^ Ly6C^intermediate (int)^ mature neutrophils. Transient neutrophil depletion (Fig. S3E) was achieved by injecting mice twice retro-orbitally (3 d before and once after ONC) and twice intraperitoneally (immediately and 7 d after ONC) with an anti-Ly6G antibody. For prolonged early neutrophil depletion (Fig. S3F), mice were injected 1 d before and once after cLI and every other day thereafter until ONC. For prolonged late neutrophil depletion (Fig. 3D), mice were injected with anti-Ly6G 1 d before and once after ONC and every other day thereafter until euthanasia, at which time blood was collected for flow analysis. Mature neutrophils were almost completely absent after immune depletion (Figs. S3G, H), yet all three strategies failed to reduce the number of regenerating axons induced by cLI (prolonged late depletion: Fig. 3E, F). Finally, we tested whether mature T cells, which play a beneficial role in several neurological diseases^24,25^, contribute to the effects of cLI. RAG1-KO mice showed the expected loss of mature T cells by CD3 staining in the retina following cLI and ONC (Figs. 3G, S3I) but this did not alter cLI-induced regeneration (Figs. 3H, I). Taken together, these results suggest that mature neutrophils and T cells are unlikely to contribute substantially to cLI-induced regeneration.

### cLI activates pro-regenerative signaling pathways in injured RGCs but does not rely on Ocm, SDF-1 or CCL5

Neuronal expression of phosphorylated signal transducer and activator of transcription 3 (pSTAT3), a marker of Jak-STAT3 pathway activation, and/or phosphorylated ribosomal protein S6 (pS6), a marker of mTOR pathway activation, is associated with a regenerative state^26,27^. cLI mice showed an elevation of both pS6 (39%) and pStat3 (47%) in RBPMS^+^ RGCs 3 days after ONC compared with mice having sham surgery (Fig. 4A-D). Because deletion of SOCS3 (suppressor of cytokine signaling-3), a negative regulator of STAT3, in RGCs promotes optic nerve regeneration^28^, we investigated whether pSTAT3 elevation might be associated with a downregulation of SOCS3. Following cLI, RGCs exhibited a small (18%) but significant decrease of SOCS3 compared to RGCs in sham-surgery controls (Fig. 4B, E). Moreover, downregulation of SOCS3 enables recombinant ciliary neurotrophic factor (CNTF) to elevate levels of pSTAT3 and to exert pro-regenerative effects^28^, raising the question of whether cLI sensitizes RGCs to CNTF. However, cLI did not enable CNTF to increase regeneration above the level induced by cLI alone (Fig. S4D-F), arguing against a role for CNTF in cLI.

**Figure 4.**
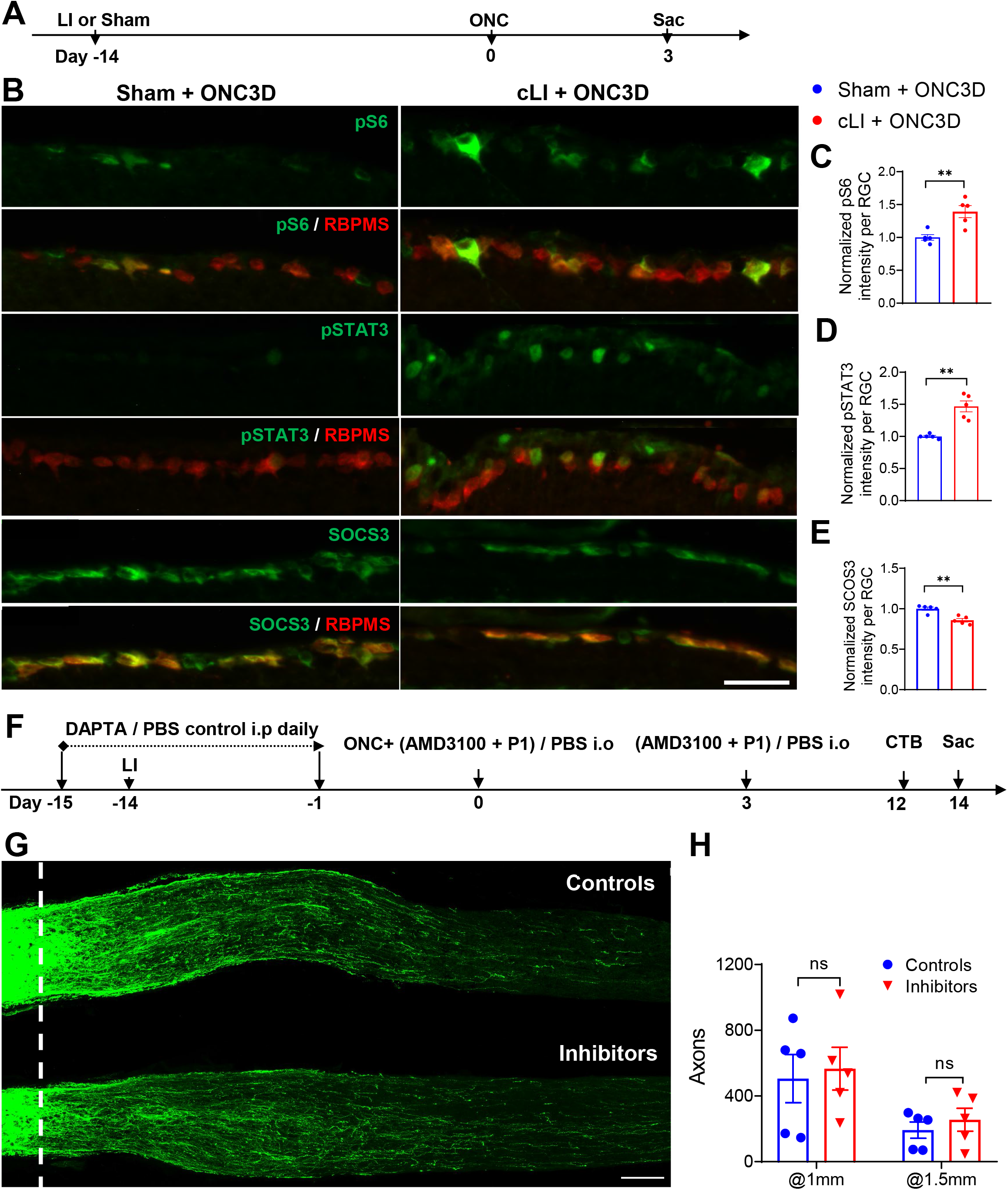
mTOR and STAT3 act as possible downstream signaling pathways for cLI induced regeneration with minimal contributions of Ocm, SDF-1, and CCL5. A. Experimental timeline. cLI or sham surgery was introduced 14 days before ONC. Mice were euthanized 3 days after ONC. B. Representative images of retinal sections showing RGCs stained with RBPMS in the ganglion cell layer co-stained with anti-pS6, anti-pSTAT3, or anti-SOCS3 antibodies. Scale bar, 50 μm. C. Quantitation of normalized pS6 immunostaining in RGCs shows increase with cLI compared to sham controls (unpaired t-test, p = 0.0047, n = 5 mice per group). D. Quantitation of normalized pSTAT3 immunostaining in RGCs in shows increase with cLI compared to sham controls (unpaired t-test, p=0.0061, n = 5 mice in each group). E. Quantitation of normalized SOCS3 intensity in RGCs shows modest decrease with cLI compared to sham controls (unpaired t-test, p = 0.0014; n = 5 mice per group). F. Experimental timeline. DAPTA or PBS was injected *i.p*. daily beginning 2 days prior to LI and continuing through one day before ONC. LI or sham surgery was introduced 14 days before ONC. AMD3100 and P1 peptide were injected intraocularly (*i.o*.) immediately and 3 days after ONC. Mice were euthanized 14 days after ONC. G. Representative longitudinal sections through the optic nerve showing similar levels of regenerating axons in mice receiving control treatments and growth factor inhibitors. *White line* indicates crush sites. Scale bar, 100 μm. H. Quantitation of regenerating axons 1 mm and 1.5 mm from crush site in (G) (unpaired t-test, p = 0.758; n = 3 or 4 mice per group).

Oncomodulin (Ocm) and stromal cell-derived factor 1 (SDF-1) are key mediators of zymosan-induced regeneration, while C-C motif chemokine receptor 5 (CCL5) mediates most of the pro-regenerative effects of virally mediated CNTF overexpression^3,29,7,9^. AMD3100, a selective antagonist to CXCR4, the primary receptor for SDF1, combined with P1 peptide, an Ocm antagonist, blocked 70% of zymosan-induced regeneration (Fig.S4A-C), while DAPTA, a selective antagonist to the CCL5 receptor CCR5, strongly suppressed regeneration induced by CNTF gene therapy^7^. However, daily intraperitoneal injection of DAPTA combined with intraocular injection of AMD3100 and P1 had no effect on regeneration induced by cLI (Fig. 4F-H). Thus, the strong pro-regenerative effect of cLI does not appear to involve the same molecular mediators as post-injury inflammation.

### Multiple LI combined with PLX treatment enables brain re-innervation

The biggest challenge of optic nerve regeneration is to re-connect RGC axons with correct brain targets for functional recovery. As shown above, multiple LI induced full-length optic nerve regeneration but without brain re-innervation (Fig. 1F, G). PLX treatment combined with cLI resulted in more axons reaching the optic chiasm (Fig. 2B, C), raising the possibility that multiple LI combined with PLX treatment (Fig. 5A) might enable brain re-innervation. Six weeks after ONC, brain visual target areas including the suprachiasmatic nucleus (SCN), lateral geniculate nucleus (LGN), and superior colliculus (SC) were examined for regenerating axons. Three out of four PLX-treated mice showed regenerating axons in the SCN though not in the LGN or SC, with the best case showing some axons entering the core region of SCN (visualized by NeuN immunostaining: Fig. 5B). None of the mice fed control chow (n=3) showed any brain innervation, as noted above. These findings demonstrate brain re-innervation can be achieved exclusively by cell non-autonomous, non-genetic manipulations.

**Figure 5.**
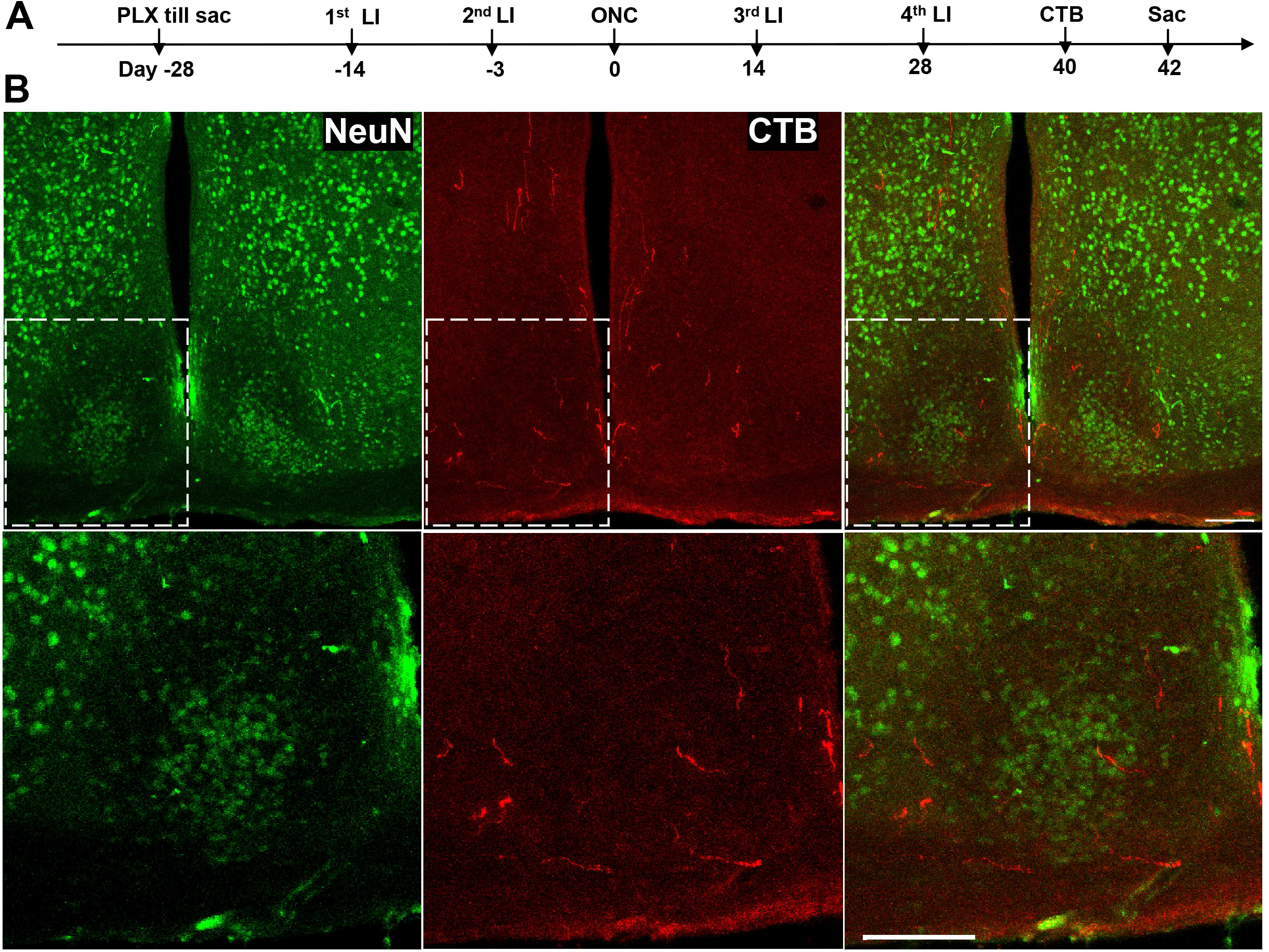
Multiple LI combined with PLX enables brain re-innervation. A. Experimental timeline. Mice received PLX treatment for 2 weeks before 1^st^ LI until euthanasia. Multiple LI was administered accordingly. Mice were euthanized 6 weeks after ONC. B. *Upper panels:* brain sections from the best case stained with anti-NeuN and anti-CTB antibodies. Lower panels display magnified images of the areas in white, dashed boxes in A, showing regenerating axons entering the core of the suprachiasmatic nucleus (SCN, visualized by NeuN immunostaining). Scale bar, 100 μm.

## Discussion

Our results show that cLI promotes far stronger optic nerve regeneration than other inflammatory manipulations studied thus far and appears to depend upon distinct cellular and molecular mechanisms. Although neutrophils and the myeloid cell-derived growth factors (Ocm, SDF1, CCL2, CNTF) that mediate the effects of post-injury pro-regenerative treatments do not seem involved, the partial effect of Ccr2 deficiency in suppressing cLI-induced regeneration in contrast to the positive effect of PLX5622 suggest that distinct subsets of reparative and detrimental peripheral/local macrophages may emerge, and/or that there may be indirect contribution from other immune cells or non-immune elements^30–32^. Combining multiple episodes of LI with PLX5622 enables some RGCs to reinnervate the suprachiasmatic nucleus within 6 weeks, a level of regeneration comparable to that induced by the most effective gene therapies or combinatorial treatments described to date^33,34^.

It should be noted that the methods used to manipulate major immune cell populations are likely to have incomplete and off-target effects, leaving remnant and rare populations such as immature neutrophils and innate lymphoid cells (ILCs)to be further explored. Also, whereas this study has focused on factors associated with inflammation, it remains possible that components of the lens *per se* contribute to regeneration, or factors derived from other tissues as a result of cLI and inflammation^35–37^. Deeper unbiased analyses to identify changes in all neuronal and non-neuronal populations, cell type-specific transcriptional changes, and ligand-receptor interactions will be essential to better understand the mechanisms underlying the phenomenon reported here^38–40^. With this new paradigm in hand, we hope that cLI can pave the way to identify more potent cell-extrinsic therapies to address currently incurable neuronal and axonal losses after CNS injury without applying genetic manipulations with potentially deleterious long-term consequences.

One final note is that the strong effects of even mild LI need to be considered in interpreting the results of studies involving intravitreal injections of viruses or other agents prior to ONC that might inadvertently injure the lens.

## Materials and methods

### Key resources table

| Reagent type (species) or resource | Designation | Source or reference | Identifiers | Additional information |
| --- | --- | --- | --- | --- |
| Strain, strain background (*Mus musculus*, male and female) | 129S1/SvImJ | The Jackson Laboratories | 002448 |  |
| Strain, strain background (*Mus musculus*, male and female) | C57BL/6J | The Jackson Laboratories | 000664 |  |
| Genetic strain, strain background (*Mus musculus*, male and female)<br>**Continued** | B6.129(Cg)-Ccr2tm2.1lfc/J | The Jackson Laboratories | 017586 |  |
| Genetic strain, strain background (*Mus musculus*, male and female) | B6.129S7-Rag1tm1Mom/J | The Jackson Laboratories | 002216 |  |
| Antibody | Rabbit anti-Cholera toxin subunit B (CTB) | GenWay Biotech | GWB-7B96E4 | IF (1:500) |
| Antibody | Rabbit polyclonal anti-beta III Tubulin | Abcam | ab18207 | IF (1:500) |
| Antibody | Rabbit polyclonal anti-Iba1 | WAKO | Cat: #019-19741 | IF (1:1000) |
| Antibody | Rat monoclonal anti-mouse CD68 | Bio-Rad | MCA1957 | IF (1:500) |
| Antibody | Anti-NeuN, Clone: A60 | Fisher Scientific | Mab377 | IF (1:200) |
| Antibody | Phospho-S6 Ribosomal Protein (Ser240/244) Rabbit mAb | Cell Signaling Technology | 5364 | IF (1:200) |
| Antibody | Guinea Pig Polyclonal anti-RBPMS | PhosphoSolutions | Cat# 1832-RBPMS | IF (1:500) |
| Antibody | Phospho-Stat3 (Tyr705) Rabbit monoclonal Ab | Cell Signaling Technology | Cat# 9145 | IF (1:200) |
| Antibody | Mouse SOCS3 Monoclonal Antibody | Fisher Scientific | MA1-19373 | IF (1:500) |
| Antibody | RFP Antibody Rabbit Polyclonal | Rockland | 600-401-379 | IF (1:500) |
| Antibody | Anti-mouse Ly6G | Bio X Cell | BP0075 | 200 µg in 200 µl (i.p.) |
| Antibody | Rat IgG2a isotype control | Bio X Cell | BE0089 | 200 µg in 200 µl (i.p.) |
| Antibody | Mouse CD3 Antibody | R&D Systems | MAB4841 | IF (1:100) |
| Antibody | Rat monoclonal anti-mouse CD16/32 (mouse Fc block) | BioLegend | 156603 | FACS |
| Commercial assay or kit | Mouse MDSC Flow Cocktail 2 with Isotype Ctrl | BioLegend | 147003 | FACS |
| Commercial assay or kit | OCT compound | Sakura Finetek | 4583 |  |
| Peptide/recombinant protein | CNTF | Pepro Tech | AF-450-50 | 0.5 µg/µl |
| Peptide/recombinant protein | CTB | Sigma | SAE0069 | 2 µg/µl |
| Peptide/recombinant protein | CTB-Alexa Fluor 555/488 Conjugate | Thermo Fisher Scientific | C-22843 /C-22841 | 2 µg/µl |
| Peptide/recombinant protein | P1 | Genscript Corp (ref. 28) | synthesized | 2.3 µg/µL |
| Peptide/recombinant protein | Bovine Serum Albumin (BSA) | Sigma-Aldrich | A9085 | 0.5 µg/µl |
| Chemical compound | DAPTA | Selleck Chemicals | S8501 | 10 µg/kg |
| Chemical compound | PLX5622 | Plexxikon Inc. |  | 1200 ppm in chow |
| Chemical compound | RBC Lysis Buffer (10X) | BioLegend | 420301 |  |
| Chemical compound | AMD3100 | Sigma | CAS 155148-31-5 | 100 µM |
| Chemical compound | Zymosan | Sigma | Z4250 | 12.5 µg/µL |
| Software, algorithm | Zen | Zeiss |  |  |
| Software, algorithm | Fiji (ImageJ) |  |  |  |
| Software, algorithm | GraphPad Prism | GraphPad | Version 8 |  |
| Other | DAPI | Sigma-Aldrich | Cat: #9542 | 1:10000 |

#### Mice

Experiments were performed at Boston Children’s Hospital with approval from the Institutional Animal Care and Use Committee. Wild type (WT) 129S1 mice (129S1/SvImJ; Strain #002448) and C57BL/6J mice (Strain #000664) of both sexes (6- to 12-week-old) were obtained from the Jackson laboratories (Bar Harbor, ME) and used in the study. CCR2^RFP/RFP^ mice [B6.129(Cg)-Ccr2tm2.1Ifc/J; Strain #017586; Jackson], and RAG1 KO mice (B6.129S7-Rag1tm1Mom/J; Strain #002216; Jackson) were originally purchased from Jackson laboratories (Bar Harbor, ME). Homozygous CCR2^RFP/RFP^ mice were crossed with C57BL/6J mice to obtain heterozygous CCR2^RFP/+^ mice. Experimental CCR2^RFP/RFP^ mice and littermate CCR2^RFP/+^ controls were generated from heterozygous CCR2^RFP/+^ breeder pairs, maintained in C57Bl/6J background, and were genotyped by Transnetyx (Cordova, TN, USA). All mice were housed under the same conditions for at least 4 days before being used in experiments. All mice were maintained in regular cages on a 12 h light/dark cycle with ad libitum access to regular food and water, except RAG1 KO mice and their matched control mice which were maintained in autoclaved cages on a 12 h light/dark cycle with ad libitum access to sterile food and water. All experiments used 6- to 12-week-old male and female 129S1 mice unless otherwise specified.

#### Mild lens injury

For surgery, mice were anesthetized by intraperitoneal injection of ketamine and xylazine. Mild lens injury was performed by puncturing the lens posterior capsule and cortex to a depth of 0.5-1 mm using a disposable 30G sharp needle while sham surgery was performed by puncturing the posterior part of the eye into the vitreous chamber with care taken not to touch the lens. Mild LI results in local response while maintaining overall transparency (shown in Fig. S1A). All eyes with mild LI look indistinguishable compared to these with sham surgery. Severe damage to the whole lens, including capsule, cortex, and nucleus indicated by global or nucleus whiteness or disintegration of the lens were excluded from the study.

#### Optic nerve crush and intravitreal injections

For surgery, mice were anesthetized by intraperitoneal injection of ketamine and xylazine. The optic nerve was intraorbitally crushed 0.5-1 mm behind the optic disc for 2-5 s using fine forceps (Dumont #5 FST), as described previously^3^. Agents to be tested were injected intravitreally in a volume of 2 μl per eye using a 33G blunt needle to avoid any lens injury. Reagents injected intravitreally include zymosan (sterilized before use); CXCR4 antagonist AMD1000; P1 peptide. CTB or CTB conjugated recombinant protein was injected intravitreally (2 μl per eye) to label regenerating axons 2 d before mice were euthanized.

#### Neutrophil depletion

To deplete neutrophils systemically, each mouse received retro-orbital or intraperitoneal (i.p.) injection of 100 μg anti-mouse Ly6G IgG (BE0075-1; Bio X Cell) or isotype-matched IgG2a (BE0089; Bio X Cell) antibody using a modified protocol^41^. To verify neutrophil depletion, blood neutrophils were evaluated by flow cytometry (BD LSRFortessa^™^, BD Biosciences) using Mouse MDSC Flow Cocktail (147003; Biolegend) and analyzed using FlowJo V10 software (Tree Star, Inc., Ashland, OR).

#### Resident macrophages depletion

To deplete resident macrophages, mice were fed chow containing 1200 ppm PLX5622 (Plexxikon Inc.) ad libitum. Control animals received the same chow (Plexxikon Inc.) but without drug. Chow consumption began 14 days prior to further experimental procedures and continued through the course of experiments to ensure sufficient and sustained depletion.

#### Immunohistochemistry and imaging

Mice were anesthetized and perfused through the heart with PBS followed by 4% PFA. Eyes and optic nerves were dissected out. Upon dissection, eyes were carefully examined under dissecting microscope for cornea damage, intraocular bleeding, severe lens damage or eye dystrophy which were used as criteria for exclusion, then post-fixed for one hour in a 4% paraformaldehyde (PFA), transferred to 30% sucrose overnight (4 °C) and embedded in Tissue-Tek (Sakura, Netherlands). Frozen sections (14 mm) were cut longitudinally on a cryostat, thaw-mounted onto glass slides (Superfrost plus, Fisher) and stored at −80 °C until further use. Antibodies are listed in key resource table. Secondary antibodies included anti-mouse, anti-rabbit, anti-guinea pig, anti-goat IgG antibodies conjugated to Alexa Fluor 488/555/594 (1: 500) (Molecular Probes, USA). Images were taken by a Nikon E800 microscope or a Zeiss LSM710 confocal microscope, then merged, cropped, and optimized using ImageJ (Fiji).

#### Quantitation of regenerating axons in the optic nerve

Axon regeneration was quantified as described previously^1^. In brief, the number of CTB-positive axons extending various distances from the injury site were counted under 400X magnification in 3–4 sections per sample. These values were normalized to the cross-sectional area of the optic nerve and extrapolated to the whole optic nerve.

#### Quantitation of RGCs in whole-mounted retina

For quantitation of surviving RGCs per mm^2^, retinal flat-mounts were stained with an antibody against βIII-tubulin. Retinas were divided into four quadrants. In each quadrant 2 independent fields were sampled, representing the center and periphery. The average number of βIII-tubulin-positive RGCs per field was determined and divided by the area of the field. Values were averaged per retina. At least 4 retinas from no fewer than 3 mice per condition were analyzed.

#### Preparation and staining of whole eye sections

Mice were perfused as described above. Eyes were collected and postfixed in 4% PFA for 1 h, transferred to 30% sucrose overnight at 4 °C, and frozen sectioned at 14 μm. Sections were incubated with primary antibodies at 4 °C overnight after blocking with appropriate sera for 1 h at RT. After washing three times, sections were incubated with the appropriate fluorescent secondary antibody and DAPI and then mounted. Primary antibodies that were used are listed in the key resource table.

#### Preparation and staining of brain sections

Mice were perfused as described above. Brains were postfixed for 48 h at 4 °C and then transferred to 30% sucrose until they sank, embedded in O.C.T., and cryostat-sectioned in the coronal plane at 50 μm. Sections were collected and stained free-floating in PBS to detect CTB-labeled growing axons and NeuN (to show the brain nucleus location), and then mounted onto slides.

#### Analysis of pS6, pSTAT3, and SOCS3 levels in RGCs

To analyze pS6, pSTAT3, and SOCS3 level in RGCs, 4 non-adjacent retinal sections from each mouse were stained simultaneously with anti-RBPMS antibodies following the steps mentioned above (see Staining of whole eye sections). To quantify the fluorescence intensity of pS6, pSTAT3, and SOCS3 in all RGCs, images acquired simultaneously with identical configurations were analyzed for each retina. At least 200 RGC per eye 3 mice each group were manually selected and measured for mean fluorescence intensity per cell using ImageJ.

#### Statistical analyses

Statistical analyses were done with GraphPad Prism 8 and the significance level was set as p < 0.05. For comparisons between two groups, two-tailed unpaired or paired t-test was used. One-way ANOVA followed by Tukey’s multiple comparison tests were performed for comparisons among three or more groups. Data are represented as mean ± SEM; P values of post hoc analyses are illustrated in the figure. n.s., not significant; *p < 0.05, **p < 0.01, ***p < 0.001, ****p < 0.0001. All details regarding statistical analyses, including the tests used, P values, exact values of n, definitions of n, are described in figure legends.

## Acknowledgments

This work was supported by the Dr. Miriam and Sheldon G. Adelson Medical Research Foundation (to L.B.), the NIH Intellectual and Developmental Disabilities Research Centers Imaging Core (HD018655) and the Flow Cytometry Core of Boston Children’s Hospital. We are grateful to Dr. Roman Giger (University of Michigan) for comments on the manuscript and to Dr. Yuqin Yin, Dr. Lili Xie, and Hui-Ya Gilbert for their critical advice and technical support. We apologize to those investigators whose important work we were unable to cite or describe owing to space constraints.

## Author contributions

**Qian Feng**: Conceptualization, Investigation, Methodology, Formal analysis, Writing - original draft, Writing - review and editing; **Kimberly Wong**: Methodology, Resources, Writing - review and editing; **Larry I. Benowitz**: Conceptualization, Investigation, Methodology, Supervision, Funding acquisition, Writing - review and editing.

---

**Figure S1.**
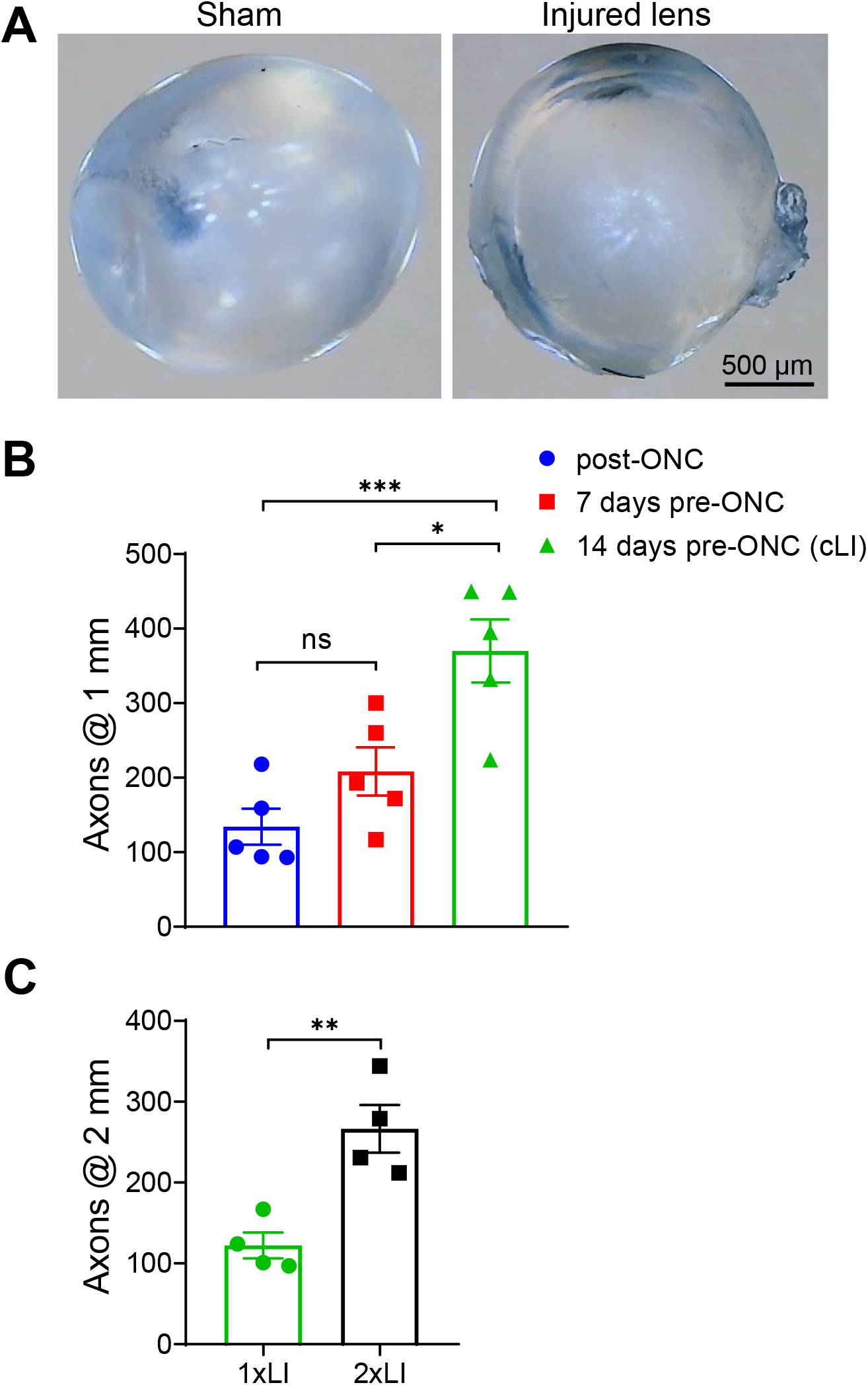
A. Representative lens images after 14 days of sham surgery or mild lens injury showing local “volcano-like” structure on the surface of the injured lens while maintaining overall transparency. Scale bar, 500 μm. B. Quantitation of regenerating axons 1 mm from crush site immediately after ONC (post-ONC), 7 days before ONC (7 days pre-ONC) and 14 days before ONC (14 days pre-ONC: cLI) (One-way ANOVA followed by Tukey’s multiple comparisons test; post-ONC vs. 7 days pre-ONC p = 0.3034; post-ONC vs. 14 days pre-ONC p = 0.0009; 7 days pre-ONC vs. 14 days pre-ONC p=0.0141; n = 3 or 4 mice per group). C. Quantitation of regenerating axons 2 mm from crush site after one (1xLI) or two LI (2x LI) (unpaired t-test, p = 0.0051; n = 3 mice per group).

**Figure S2.**
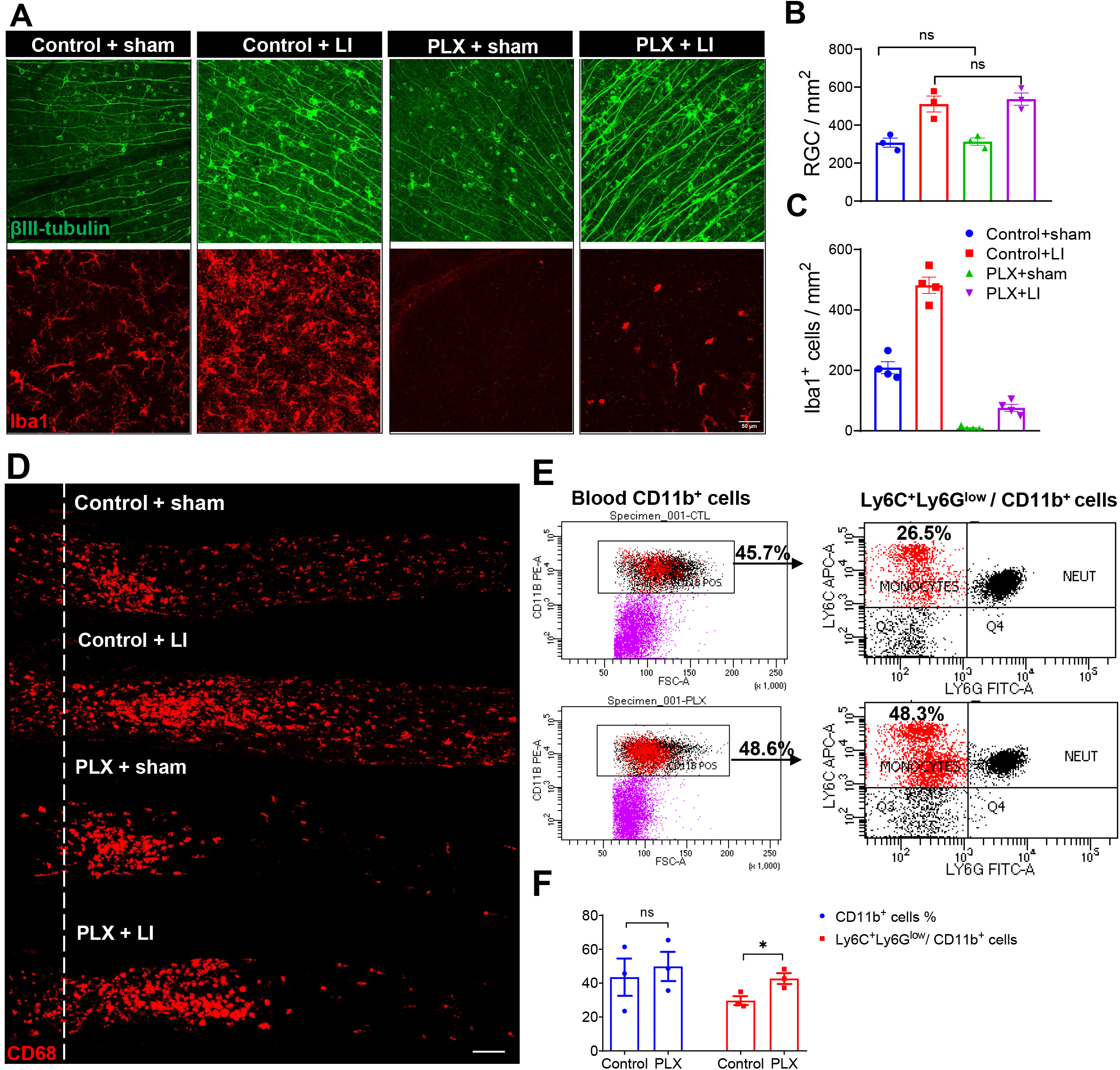
A. Representative whole-mounted retinas showing βIII-tubulin labeled RGCs and Iba1 labeled macrophages 4 weeks after ONC. Scale bar, 50 μm. B. Quantitation of βIII-tubulin+ cells for surviving RGCs in (A) (one-way ANOVA followed by Tukey’s multiple comparisons test; Control + Sham vs. PLX + Sham, p= 0.9993; Control + LI vs. PLX + LI, p= 0.9316; n=3 mice each group). C. Quantitation of Iba1^+^ cells in (A) (Mean numbers are 209 in Control + sham vs. 481 in Control + LI vs. 5 in PLX + sham vs. 66 in PLX + cLI). D. Representative longitudinal sections through the optic nerve showing CD68^+^ cells. A population of CD68^+^ cells persisted in the optic nerve especially around crush sites after PLX treatment combined with sham surgery or cLI. *White line:* crush site. Scale bar, 100 μm. E. Flow cytometric analysis of blood collected from mice receiving PLX chow or control chow after 4 weeks. Dot-plot of monocytes (CD11b^+^ Ly6C^+^Ly6G^low^) in blood. The percentages of CD11 b^+^ cells and monocytes in CD11 b^+^ cells are shown. F. Quantitation of the percentages of CD11 b^+^cells and monocytes in CD11b^+^ cells in (D) (unpaired t-test; CD11b^+^ cell %: control vs. PLX, p = 0.6739; monocytes / CD11b^+^ cells: control vs. PLX, p = 0.0349; n = 3 mice in each group).

**Figure S3.**
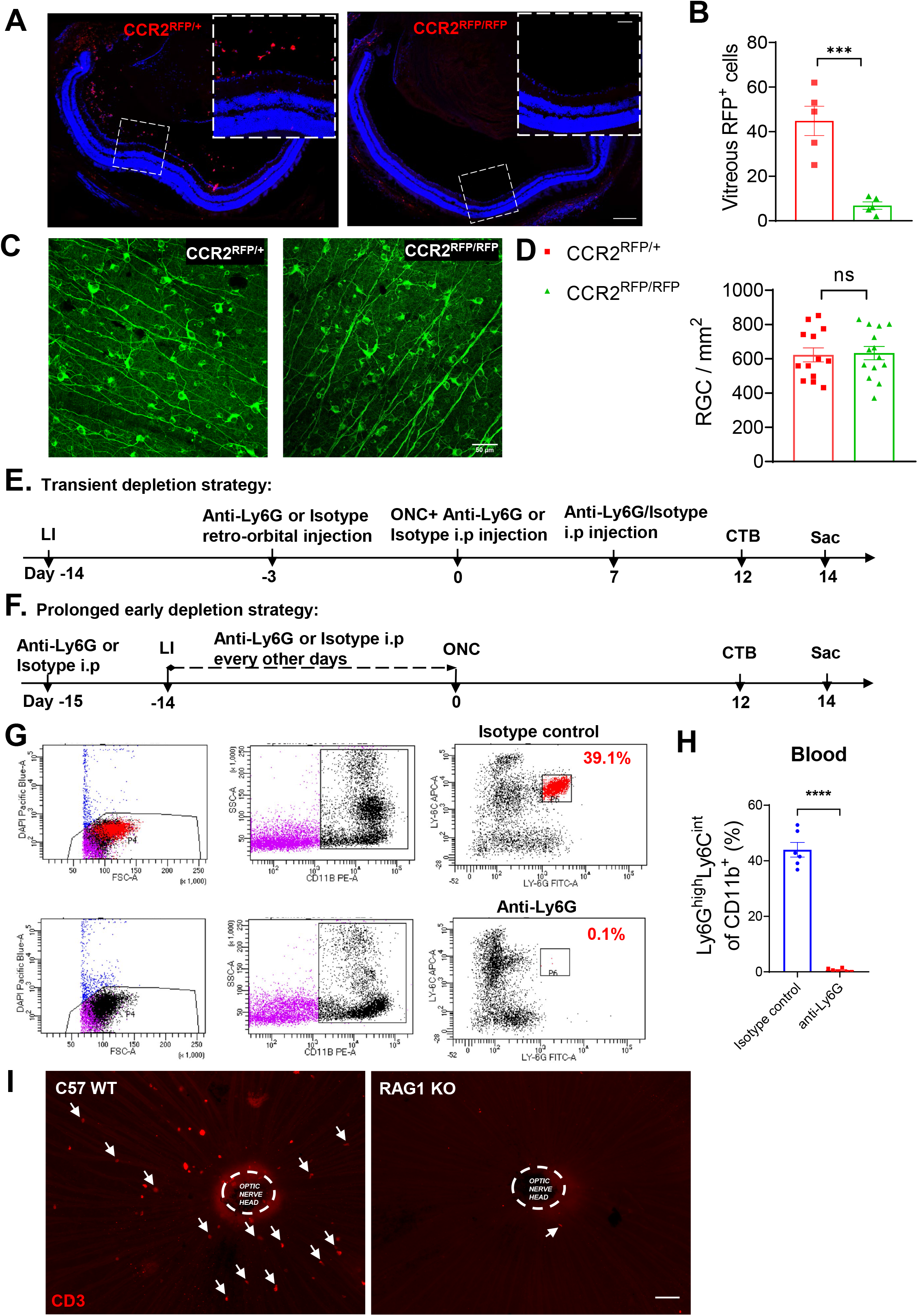
A. Representative whole eye sections showing much less RFP^+^ cells in the vitreous chamber in CCR2^RFP/RFP^ mice than CCR2^RFP/+^mice. The white dashed boxes in the right upper are the magnified images showing RFP^+^ cells. Scale bar, 200 μm; Enlarged scale bar, 50 μm. B. Quantitation of vitreous RFP^+^ cell number in (A) (unpaired t-test, p = 0.0005; n = 5 mice in each group; at least 4 non-sequential sections were analyzed for each mouse). C. Representative whole-mounted retinas showing βIII-tubulin labeled RGCs in CCR2^RFP/RFP^ mice and CCR2^RFP/+^mice. Scale bar, 50 μm. D. Quantitation of βIII-tubulin^+^ RGCs in (C) (unpaired t-test, p= 0.853; n = 6 or 7 mice in each group). E. Experimental timeline for transient depletion strategy. F. Experimental timeline for prolonged early depletion strategy. G. Flow cytometric analysis of blood collected from mice receiving isotype control injections or anti-Ly6G injections. Dotplot of mature neutrophils (CD11b^+^ Ly6G^high^Ly6C^int^) in blood 14 days post-injections using prolonged late depletion strategy (Fig. 3D). The percentages of neutrophils in CD11b^+^ cells were shown. H. Quantitation of the percentages of mature neutrophils in CD11b^+^ cells in (G) (unpaired t-test, p < 0.0001; n = 6 mice in each group). I. Representative whole-mounted retinas showing lack of CD3^+^ cells near the optic nerve head in RAG1 KO mice compared to C57 WT mice. *White arrow:* CD3^+^cell. Scale bar, 50 μm.

**Figure S4.**
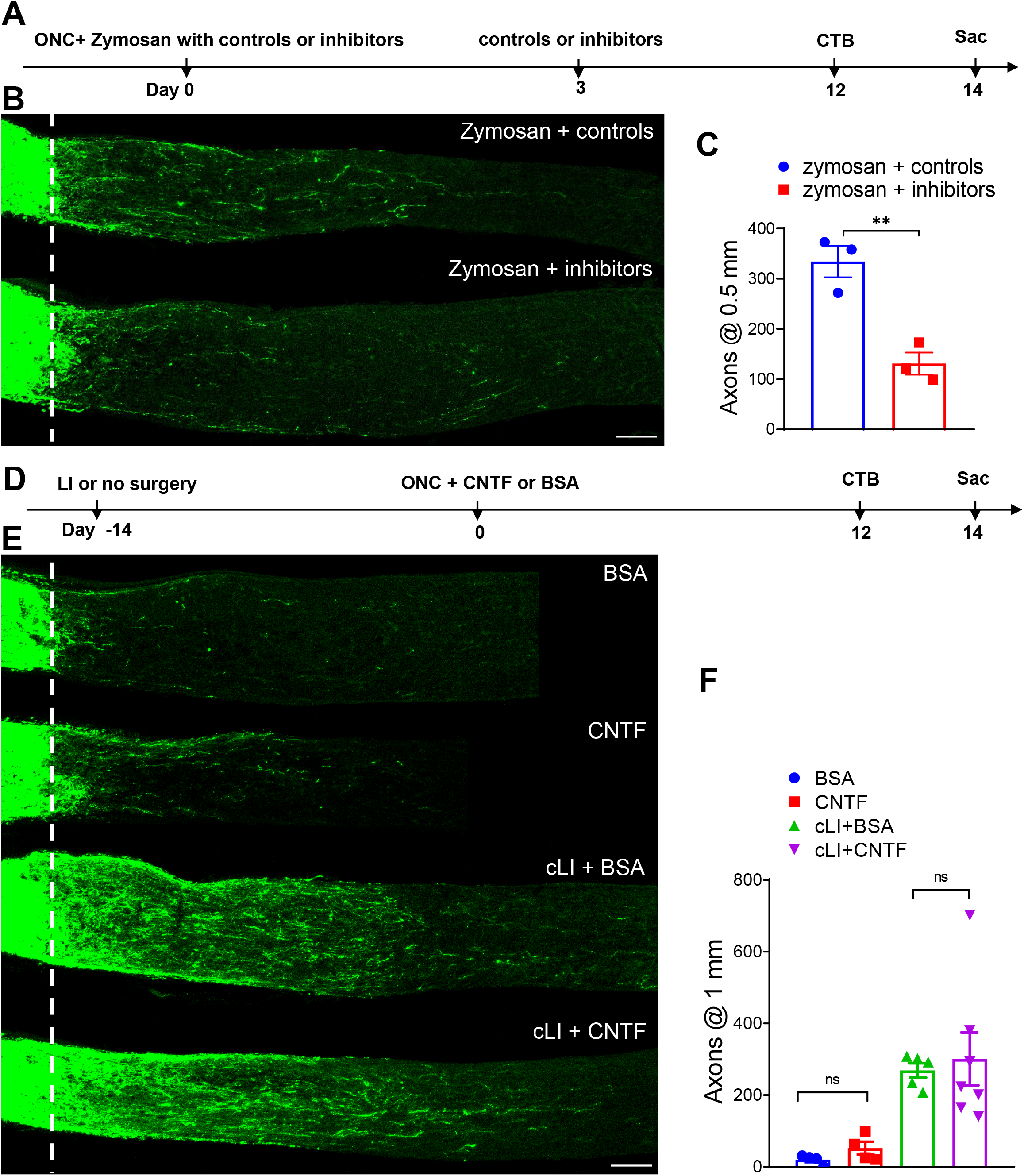
A. Experimental timeline. Zymosan with control peptide or with AMD3100 and P1 peptide were injected intraocularly (*i.o*.) immediately after and 3 days after ONC. Mice were euthanized 14 days after ONC. B. Representative longitudinal sections through the optic nerve showing much fewer regenerating axons in mice receiving zymosan plus inhibitors than in mice receiving zymosan plus controls. *White line:* crush site. Scale bar, 100 μm. C. Quantitation of regenerating axons at 0.5 mm from crush sites in (C) (unpaired t-test, 0.0061; n = 3 mice per group). D. Experimental timeline. LI (or no surgery) was introduced 14 days before ONC, and CNTF or BSA were injected intraocularly (i.o) immediately after ONC. Mice were euthanized 14 days later. E. Longitudinal sections through the optic nerve showing regenerating axons 14 days post-ONC. CNTF injection alone post-ONC resulted in few regenerating axons, and cLI combined with CNTF (cLI + CNTF) showed similar levels of regeneration compared to cLI combined with BSA (cLI + BSA). *White line* indicates crush sites. Scale bar, 100 μm. F. Quantitation of regenerating axons 1 mm from crush sites in (C) (unpaired t-test, BSA vs. CNTF, p = 0.152; cLI + BSA vs. cLI + CNTF, p = 0.73; n = 4 or 5 mice each group).

## References

1. Leon, S., Yin, Y., Nguyen, J., Irwin, N. & Benowitz, L. I. Lens Injury Stimulates Axon Regeneration in the Mature Rat Optic Nerve. J. Neurosci. 20, 4615–4626 (2000).

2. Yin, Y. et al. Macrophage-Derived Factors Stimulate Optic Nerve Regeneration. J. Neurosci. 23, 2284–2293 (2003).

3. Yin, Y. et al. Oncomodulin is a macrophage-derived signal for axon regeneration in retinal ganglion cells. Nat. Neurosci. 9, 843–852 (2006).

4. Kurimoto, T. et al. Neutrophils Express Oncomodulin and Promote Optic Nerve Regeneration. J. Neurosci. 33, 14816–14824 (2013).

5. Baldwin, K. T., Carbajal, K. S., Segal, B. M. & Giger, R. J. Neuroinflammation triggered by β-glucan/dectin-1 signaling enables CNS axon regeneration. Proc. Natl. Acad. Sci. 112, 2581–2586 (2015).

6. Sas, A. R. et al. A new neutrophil subset promotes CNS neuron survival and axon regeneration. Nat. Immunol. 21, 1496–1505 (2020).

7. Xie, L., Yin, Y. & Benowitz, L. Chemokine CCL5 promotes robust optic nerve regeneration and mediates many of the effects of CNTF gene therapy. Proc. Natl. Acad. Sci. 118, e2017282118 (2021).

8. Liu, Y.-F. et al. CXCL5/CXCR2 modulates inflammation-mediated neural repair after optic nerve injury. Exp. Neurol. 341, 113711 (2021).

9. Xie, L. et al. Monocyte-derived SDF1 supports optic nerve regeneration and alters retinal ganglion cells’ response to Pten deletion. Proc. Natl. Acad. Sci. U. S. A. 119, e2113751119 (2022).

10. Garcia-Bonilla, L. et al. Endogenous Protection from Ischemic Brain Injury by Preconditioned Monocytes. J. Neurosci. Off. J. Soc. Neurosci. 38, 6722–6736 (2018).

11. Pietrucha-Dutczak, M., Amadio, M., Govoni, S., Lewin-Kowalik, J. & Smedowski, A. The Role of Endogenous Neuroprotective Mechanisms in the Prevention of Retinal Ganglion Cells Degeneration. Front. Neurosci. 12, (2018).

12. Gidday, J. M. Adaptive Plasticity in the Retina: Protection Against Acute Injury and Neurodegenerative Disease by Conditioning Stimuli. 26 (2019).

13. Kim, E. & Cho, S. CNS and peripheral immunity in cerebral ischemia: partition and interaction. Exp. Neurol. 335, 113508 (2021).

14. Lu, X. & Richardson, P. M. Inflammation near the nerve cell body enhances axonal regeneration. J. Neurosci. 11, 972–978 (1991).

15. Neumann, S. & Woolf, C. J. Regeneration of Dorsal Column Fibers into and beyond the Lesion Site following Adult Spinal Cord Injury. Neuron 23, 83–91 (1999).

16. Kwon, M. J. et al. Contribution of macrophages to enhanced regenerative capacity of dorsal root ganglia sensory neurons by conditioning injury. J. Neurosci. Off. J. Soc. Neurosci. 33, 15095–15108 (2013).

17. Kwon, M. J. et al. CCL2 Mediates Neuron-Macrophage Interactions to Drive Proregenerative Macrophage Activation Following Preconditioning Injury. J. Neurosci. Off. J. Soc. Neurosci. 35, 15934–15947 (2015).

18. Niemi, J. P., DeFrancesco-Lisowitz, A., Cregg, J. M., Howarth, M. & Zigmond, R. E. Overexpression of the monocyte chemokine CCL2 in dorsal root ganglion neurons causes a conditioning-like increase in neurite outgrowth and does so via a STAT3 dependent mechanism. Exp. Neurol. 275 Pt 1, 25–37 (2016).

19. Niemi, J. P. et al. The Conditioning Lesion Response in Dorsal Root Ganglion Neurons Is Inhibited in Oncomodulin Knock-Out Mice. eNeuro 9, ENEURO.0477-21.2022 (2022).

20. Yang, X., Liu, R., Xu, Y., Ma, X. & Zhou, B. The Mechanisms of Peripheral Nerve Preconditioning Injury on Promoting Axonal Regeneration. Neural Plast. 2021, (2021).

21. Hilla, A. M., Diekmann, H. & Fischer, D. Microglia Are Irrelevant for Neuronal Degeneration and Axon Regeneration after Acute Injury. J. Neurosci. 37, 6113–6124 (2017).

22. Lei, F. et al. CSF1R inhibition by a small-molecule inhibitor is not microglia specific; affecting hematopoiesis and the function of macrophages. Proc. Natl. Acad. Sci. 117, 23336–23338 (2020).

23. Fujimura, N. et al. CCR2 inhibition sequesters multiple subsets of leukocytes in the bone marrow. Sci. Rep. 5, 11664 (2015).

24. Walsh et al. MHCII-independent CD4+ T cells protect injured CNS neurons via IL-4. J. Clin. Invest. (2015) doi:10.1172/jci76210.

25. Shi, L. et al. Treg cell-derived osteopontin promotes microglia-mediated white matter repair after ischemic stroke. Immunity (2021) doi:10.1016/j.immuni.2021.04.022.

26. Park, K. K. et al. Promoting Axon Regeneration in the Adult CNS by Modulation of the PTEN/mTOR Pathway. Science 322, 963–966 (2008).

27. Luo, X. et al. Enhanced Transcriptional Activity and Mitochondrial Localization of STAT3 Co-induce Axon Regrowth in the Adult Central Nervous System. Cell Rep. 15, 398–410 (2016).

28. Smith, P. D. et al. SOCS3 Deletion Promotes Optic Nerve Regeneration In Vivo. Neuron 64, 617–623 (2009).

29. Yin, Y. et al. Oncomodulin links inflammation to optic nerve regeneration. Proc. Natl. Acad. Sci. 106, 19587–19592 (2009).

30. Li, Y. et al. Delayed microglial depletion after spinal cord injury reduces chronic inflammation and neurodegeneration in the brain and improves neurological recovery in male mice. Theranostics 10, 11376–11403 (2020).

31. Li, Z. et al. M-CSF, IL-6, and TGF-β promote generation of a new subset of tissue repair macrophage for traumatic brain injury recovery. Sci. Adv. 7, eabb6260 (2021).

32. Babaeijandaghi, F. et al. Metabolic reprogramming of skeletal muscle by resident macrophages points to CSF1R inhibitors as muscular dystrophy therapeutics. Sci. Transl. Med. 14, eabg7504 (2022).

33. Yang, S.-G. et al. Strategies to Promote Long-Distance Optic Nerve Regeneration. Front. Cell. Neurosci. 14, 119 (2020).

34. Williams, P. R., Benowitz, L. I., Goldberg, J. L. & He, Z. Axon Regeneration in the Mammalian Optic Nerve. Annu. Rev. Vis. Sci. 6, 195–213 (2020).

35. Fischer, D., Hauk, T. G., Müller, A. & Thanos, S. Crystallins of the β/γ-superfamily mimic the effects of lens injury and promote axon regeneration. Mol. Cell. Neurosci. 37, 471–479 (2008).

36. Li, Y. et al. Macrophage recruitment in immune-privileged lens during capsule repair, necrotic fiber removal, and fibrosis. iScience 24, 102533 (2021).

37. Menko, A. S. et al. Resident immune cells of the avascular lens: Mediators of the injury and fibrotic response of the lens. FASEB J. 35, e21341 (2021).

38. Liu, K. & Geng, Y. Exploring Lens-Injury-Derived Effect on Optic Nerve Regeneration after Optic Nerve Lesion with Cataractogenic Lens Injury by the RNA-Seq Profiling. SSRN Scholarly Paper at https://doi.org/10.2139/ssrn.3421584 (2019).

39. Benhar, I. et al. Temporal single cell atlas of non-neuronal retinal cells reveals dynamic, coordinated multicellular responses to central nervous system injury. 2022.07.10.499469 Preprint at https://doi.org/10.1101/2022.07.10.499469 (2022).

40. Salvador, A. F. M. & Kipnis, J. Immune response after central nervous system injury. Semin. Immunol. 101629 (2022) doi:10.1016/j.smim.2022.101629.

41. Kang, L. et al. Neutrophil extracellular traps released by neutrophils impair revascularization and vascular remodeling after stroke. Nat. Commun. 11, 2488 (2020).

